# Environmental drivers of soil virus diversity across UK habitats

**DOI:** 10.64898/2026.10.20.760436

**Authors:** Sean Meaden, Briony Jones, Tim Goodall, Alfie Sloman, Susheel Bhanu Busi, Rob Griffiths

## Abstract

Soil viruses are critical members of the soil microbiome, shaping terrestrial ecosystem processes and microbial dynamics. However, the environmental factors governing their landscape-scale biogeography remain poorly characterised. Here, we analysed bulk soil metagenomic data from the UKCEH Countryside Survey across broad habitat classes to identify the key drivers of viral diversity and community assembly in UK soils. We show that soil pH is the primary predictor of both viral species richness and overall community composition, alongside significant contributions from the co-related soil organic matter and broad habitat type. Viral species diversity exhibited a strongly non-linear relationship with pH, peaking in neutral to slightly acidic soils. Viral community dissimilarity strongly correlated with 16S rRNA bacterial community structure across all taxonomic ranks, with the strongest association at ASV-level resolution. Despite high spatial turnover among viral contigs, and significant habitat-specific enrichment of particular anti-phage defence systems, viral lifestyle proxies like integrase gene frequencies showed no habitat-specific variation. Together, our results demonstrate that UK soil viral communities are primarily structured by soil pH and habitat parameters and closely coupled to patterns of bacterial host distribution, providing a baseline for future investigations and monitoring of change.

## Introduction

The viral composition of soil microbiomes has been largely overlooked despite obvious importance for regulating host populations, in turn impacting ecosystem function (Jansson and Wu, 2022). The viruses of bacteria, phages, make up the majority of soil viruses (Emerson, 2019), contributing to carbon cycling via bacterial cell lysis and through viral biomass contributing to the soil carbon pool (Jansson and Wu, 2022). They also shape bacterial physiology through short term manipulation of metabolism during infection, known as virocells (Braga et al. 2021), or from longer term associations like lysogeny. In these cases, they may contribute to soil function via modification of host activity, providing transient fitness benefits to their hosts.

However, we lack a complete understanding of the major drivers of viral community composition and diversity. Temperate soils, including the UK, hold the highest levels of belowground carbon (Crowther et al. 2019). Warming temperatures are predicted to increase bacterial necromass, as bacteria typically outcompete fungi at higher temperatures (Hu et al. 2023). The implications for soil viruses are less clear as their fate depends on that of the host, as well as the extracellular conditions. A direct effect of warming is that drier soils typically reduce extracellular viral densities (Santos-Medellín et al. 2023), while an indirect effect is that higher temperatures will increase bacterial abundances, in turn selecting for lytic infections (Jansson and Wu, 2023). These are complex interactions that could vary among different soil types. Therefore, a clear baseline encompassing a range of soil types is needed in order to predict the effects of future climate scenarios.

Here we leveraged soil metagenomic data collected as part of a long-term UK soil monitoring programme to assess viral biodiversity. We compared viral sequences from broad habitat classes that span a range of soil pH levels. We integrate co-located environmental measurements to identify underlying drivers of diversity, community assembly and prevalence across the UK. Ultimately this work aims to contextualise the UK soil virus community and provide a starting-point for interpreting future patterns of viral biogeography and ecology.

## Methods

### Survey details and soil collection

Soil samples were collected as part of the UKCEH Countryside Survey, as described by Jones, Goodall et al. (2021). The Countryside Survey is a long-term monitoring programme of the UK landscape that began in 1978. Sampling locations comprise randomly selected 1-km squares stratified to represent the major environmental classes occurring across Great Britain. Soil sampling is undertaken alongside vegetation and other environmental surveys.

Briefly, topsoil cores (15 cm depth, 5 cm diameter) were collected and transported to the laboratory on the day of sampling. Physicochemical analyses included soil pH, moisture content, organic matter (determined by loss on ignition), and total carbon and nitrogen concentrations. Prior to DNA extraction, subsamples from beneath the root-dense organic horizon were isolated, homogenised and stored at −20 °C.

For the present study, a subset of soils was selected to encompass a broad gradient in soil pH and organic matter content. Sampling squares were selected across the national soil pH gradient, and two contrasting soils were chosen from each square, representing relatively high- and low-intensity land management. This paired design generated within-square contrasts in soil organic matter, with more intensively managed soils generally having lower organic matter content and less intensively managed soils having higher organic matter content. An additional 12 soil samples were included from a separate sampling exercise originally designed to contrast arable and grassland management across two contrasting parent materials. These samples were included to extend the environmental range represented in the present survey though the factorial structure of their original sampling design was not considered in subsequent analyses. The resulting dataset therefore captured broad variation in soil pH among locations while incorporating local contrasts in organic matter associated with land management.

### Metagenome sequencing, processing and co-assembly

DNA was extracted from 5 g of each frozen, archived soil sample using the DNeasy PowerMax Soil Kit (QIAGEN), following the manufacturer’s instructions. Extracted DNA was submitted to a commercial sequencing provider for quality assessment, library preparation and shotgun metagenomic sequencing. Following library quality control, libraries were sequenced on an Illumina NovaSeq platform, generating 150 bp paired-end reads (2 × 150 bp). Read quality was assessed using FastQC and MultiQC, Illumina Universal adaptor sequences (AGATCGGAAGAGC and AGATCGGAAGAGC) were removed using Cutadapt 4.1 (Martin, 2011). Reads were quality trimmed using Sickle Paired End (Joshi and Fass, 2011) with a sliding window 0.1 times read length and a minimum Phred quality score of 20. Reads less than 20bp post trimming were discarded.

Trimmed reads from all 100 samples were co-assembled using MEGAHIT v1.2.9 across 120 threads (Li et al., 2015). K-mer sizes were informed by the “meta-large” preset but modified based on the dataset size and available computational capacity in order to reduce assembly graph complexity (k-min 37, k-max 137, k-step 10). Contigs shorter than 1000bp were removed. Read data are available from accession PRJEB75011.

### Viral Database Curation

Viral sequences were classified by processing the complete co-assembly with the GeNomad end-to-end pipeline (version 1.11.0, database version 1.9, Camargo et al. 2024). Due to the size of the input assembly, this was processed as sub-files of 100k sequences (and one file of 24572 sequences). The outputs were then combined into a master dataset. This initial analysis yielded an output of 79,560 viral sequences. From this initial database, two distinct viral databases were made. Firstly, we curated a ‘stringent’ database of 3,082 viral sequences using the criteria described in Graham et al. (2024). Namely these sequences had a minimum length of 5kb and for sequences between 5-10kb a virus score > 0.9 was required, a marker enrichment score > 2, and at least one viral marker gene. For sequences > 10kb a virus score > 0.8 was required with at least one marker gene. We then curated a ‘relaxed stringency’ database of 58,703 viral sequences by filtering to include only sequences with a minimum length of 1kb that also contained at least one viral hallmark gene, with the aim of capturing viral genome fragments, rather than complete, or semi-complete, genomes.

Sequences from each database were then clustered at 95% ANI and 85% alignment fraction of the shorter sequence, following the approach of Nayfach et al. (2021), to remove redundant sequences. We note this process only removed 200 sequences from the ‘relaxed’ database, and none from the ‘stringent’ database, presumably due to co-assembly of samples collapsing highly similar sequences. We refer to these filtered sequences as vOTUs herein, although definitions vary in the literature.

### vOTU Table Generation

Trimmed reads were then mapped to both the ’relaxed’ and ’stringent’ viral contig databases using bowtie2 in ’sensitive’ mode. CoverM was used to collect mapping statistics, including mean coverage, count of mapped reads, reads recruited per base, and contig lengths. A minimum read percent identity of 95% and a minimum read alignment length of 75% were required. This resulted in two vOTU abundance tables, which we took forward for ecological analyses.

We compared the number of reads recruited for each sample for both the ‘relaxed’ and ‘stringent’ classification vOTU datasets. A strong linear relationship was observed between the two datasets viral read recruitment (R^2^ = 0.86). Therefore, we decided to move forward with the ’relaxed’ classification dataset for viral community ecology analysis, using the ‘vegan’ package in R (v 2.7-3). To assess how representative our sequencing was of the overall community we used coverage analysis implemented in Nonpareil3 (Rodriguez-R et al, 2018, version 3.5.5). For each sample, k-mer based coverage was estimated at various sequencing depths, using 100 subsets per sample. Mean redundancy is plotted for each environment in Fig.S1. Overall, samples had a median redundancy of 0.41 when using their full sequence datasets.

Rarefaction curves were also plotted across the various habitats to assess how much of the viral community we had recovered, using the ‘relaxed’ database. While many samples did reach an asymptote this was not universal, however we observed no differences among habitat types regarding the number of samples that reached (or neared) sampling saturation (Fig. S2). We note that these values, due to the short length of many contigs, represent saturation of viral contig sequences, and don’t necessarily reflect the number of viral species per environment.

### Diversity analyses

Alpha diversity metrics (species richness and Shannon’s index) were modelled against the soil metadata to identify predictors of viral diversity. These metadata variables consisted of pH, organic matter (loss on ignition), broad habitat class, and parent material (underlying bedrock). We used general additive models (GAM) due to the strongly non-linear effect of pH. Because total sequencing reads varied across samples, we included the natural logarithm of the sequencing library size as an offset term in a negative binomial GAM. This standardises the expected richness relative to sequencing effort, controlling for uneven depth without requiring rarefaction or data transformation, although models lacking this offset parameter gave qualitatively identical results. To confirm that sequencing depth was not systematically confounded by habitat type prior to modelling, quality-filtered read counts per sample were compared across broad habitat classes using a linear model. Residuals were inspected to confirm normality, and pairwise post-hoc comparisons were conducted using the ‘emmeans’ package with Tukey’s HSD adjustment. Sequencing depth did not differ significantly across habitats (p > 0.05; Fig. S3, Table S1).

The Countryside Survey included a paired sampling strategy with proximally co-located (1km^2^) high and low land-use intensities. Each pair includes a high and low land use intensity, although by definition they are no longer the same broad habitat class. Initial modelling included a mixed-effects negative binomial regression with ’Pair’ as a random intercept to account for the paired experimental design. However, the variance explained by the pairing was estimated to be zero (yielding a singular fit boundary), indicating that the paired design did not account for any meaningful variation in viral richness, we therefore removed this variable from our alpha-diversity analyses. In essence, the viral communities of these paired samples are as different to each other as they are to the other sampling locations.

For community composition analysis, vOTU tables were converted to relative abundances. Beta diversity was evaluated using Non-Metric Multidimensional Scaling (NMDS) based on Bray-Curtis dissimilarity matrices. A smooth response surface was fitted to the ordination for soil pH was fitted using generalized additive models (ordisurf). To quantify the variance explained by environmental drivers (pH, loss on ignition, broad habitat class, parent material and intensity), we conducted a Permutational Multivariate Analysis of Variance (PERMANOVA; adonis2) using Bray-Curtis distances with 999 permutations.

### Indicator analysis

To identify viral contigs characteristic of each broad habitat class, we conducted Indicator Species Analysis using the ‘indicspecies’ package in R. Data were first transformed to relative abundances. Group-site associations were evaluated using the group-size-corrected Indicator Value index restricted to individual habitat types with 999 permutations. Habitat classes represented by fewer than four samples were excluded from analysis (Bracken and Dwarf Shrub heathland). To project indicator taxa onto community structure, vOTUs displaying significant habitat associations (p < 0.05) were fitted as vectors onto a Bray-Curtis NMDS ordination using ‘envfit’ (100 permutations), with vector lengths scaled proportionally to their indicator statistic.

### Bacterial community analysis

To assess co-variance between viral and bacterial community beta-diversity, Mantel tests were conducted comparing viral communities against a 16S rRNA dataset generated from the same soil cores. Viral read counts were converted to relative abundance and pairwise Bray-Curtis dissimilarity matrices were calculated. Similar dissimilarity matrices were calculated for bacterial communities at six taxonomic ranks (ASV, genus, family, order, class, and phylum). Mantel correlations between viral beta-diversity and bacterial beta-diversity matrices, at each phylogenetic level, were evaluated using Spearman’s rank correlation with 9999 permutations.

### Functional trait analysis

We selected viral sequences scored as medium or high quality, or complete, based on CheckV predictions. We passed these through the DRAM2 pipeline (version 2.0.0beta) using the preformatted KOfam HMM profiles provided with DRAM2 (accessed Nov.2025). The pipeline was implemented as a Nextflow (v25.10.2) workflow using apptainer. Contig-level read counts from the previous analyses were linked to the KOfam IDs by inner joining both datasets on contig names. Contig read counts were then aggregated by KEGG Orthology identifier (kofam_id) per sample to generate a primary KO-level count matrix, retaining features with >10 reads in at least three samples.

One-versus-all differential abundance testing across habitat types was performed using DESeq2 with local dispersion fitting and library size factor normalization (i.e., sum of the read counts per sample). Significantly enriched features were mapped to KEGG pathways retrieved from the KEGG REST API (KEGGREST::keggLink). Log2-fold changes were averaged across identical functional descriptions within each soil type to summarise differential functional enrichment. Annotations were also searched for integrases and AMR genes from the GeNomad pipeline analysis.

### Defence system analysis

To identify relationships with defence systems that target phages, we screened the original metagenomic assembly for anti-phage systems. We first removed sequences shorter than 1kb, then divided the assembly into 200 subsets for computational tractability. We then ran the PADLOC tool to identify defence systems in the contigs, with the underlying prodigal annotations set to metagenomic mode. We then combined the results, extracted contigs carrying defence systems and mapped the reads from each sample to this database. CoverM was used to generate coverage read recruitment statistics for each of these contigs. Almost all contigs carried a single defence system (n = 90,208, median = 1.01) due to the fragmented nature of the assembly. An indicator analysis was carried out to identify the specificity and fidelity of each defence system type across the habitat classes.

## Results

### Viral community ecology

We aimed to describe the viral diversity present in UK soils across diverse land use types, using metagenomic data collected as part of an ongoing national monitoring programme. We first mined the assembled metagenome data for viral sequences and collated this into a database of viral sequences. Of the measured environmental parameters, we found a strongly non-linear effect of pH (χ^2^ (67) = 28.7, p < 0.0001), with intermediate soil pH values showing the highest viral diversity (Fig.1). We also found a significant effect of organic matter (χ^2^ (67) = 6.1, p < 0.01) and the broad habitat class from where samples were collected (χ^2^ (67) = 23, p < 0.01). We found no significant effect of parent material (χ^2^ (67) = 19, p = 0.31) (Fig. 1. Regarding pH, vOTU alpha diversity was highest in improved grasslands (average pH 5.83) with reduced diversity in both slightly alkaline soils (calcareous grasslands, pH 7.7) and acidic soils (dwarf shrub heath, pH 3.71) (Fig.1).

**Fig. 1.**
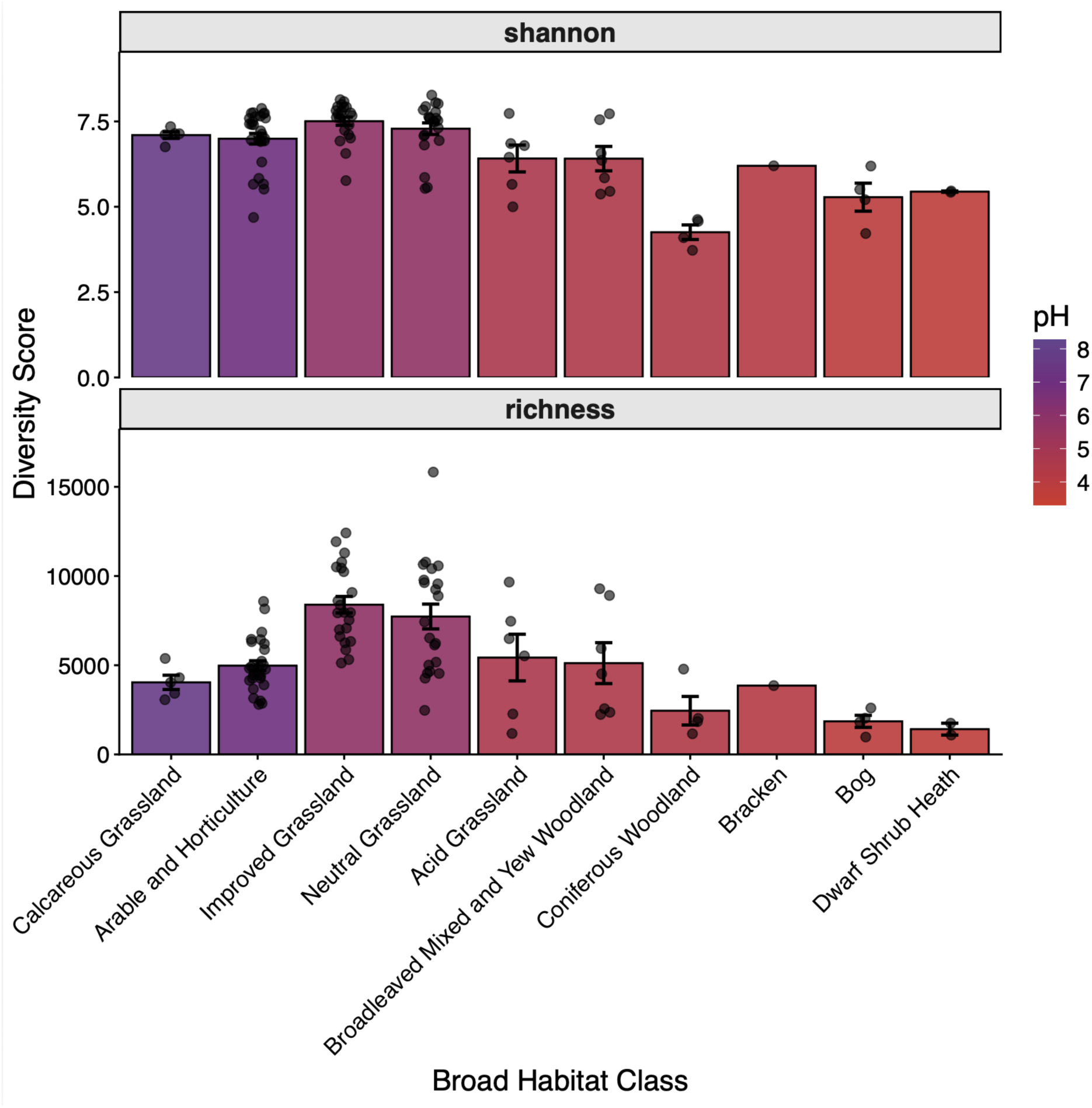
pH and land use variation in soil viral diversity. Bulk-metagenome derived viral sequences vary in species richness (bottom) and Shannon diversity (top) across a gradient of pH. Improved grasslands show maximum diversity with a mean pH of 5.83. Lower diversity is observed in both slightly alkaline soils (calcareous grasslands, pH 7.7) and acidic soils (dwarf shrub heath, pH 3.71). Error bars show 1 SE of the mean pH.

Interestingly, although we saw similar trends with bacterial species richness there were some differences. Using a dataset of 16S ASVs generated from the same soil samples we assessed species richness across the habitats. In this case, arable and horticulture soil samples showed the highest species richness, with a mean pH of 6.95. As with the viral dataset, alpha diversity was reduced in the more acidic soils and the calcareous grassland samples (Fig. S4).

Although sites in the Countryside Survey were sampled in spatially proximal pairs of contrasting land-use intensities, accounting for pairing as a random effect yielded zero variance in the model. This suggests that soil pH and habitat classes, rather than any geographic proximity, are the main drivers of viral richness. The exact sampling locations are not available to preserve landowner privacy, meaning we were unable to assess biogeographic trends further.

Similarly to the alpha diversity analysis, the same environmental parameters were significant predictors of viral community composition, with primary material also being a significant predictor (Table 1). Overall, pH was the major driver of viral community composition (Fig. 2A), although many of these variables, including pH, covary with habitat type by definition.

**Fig. 2.**
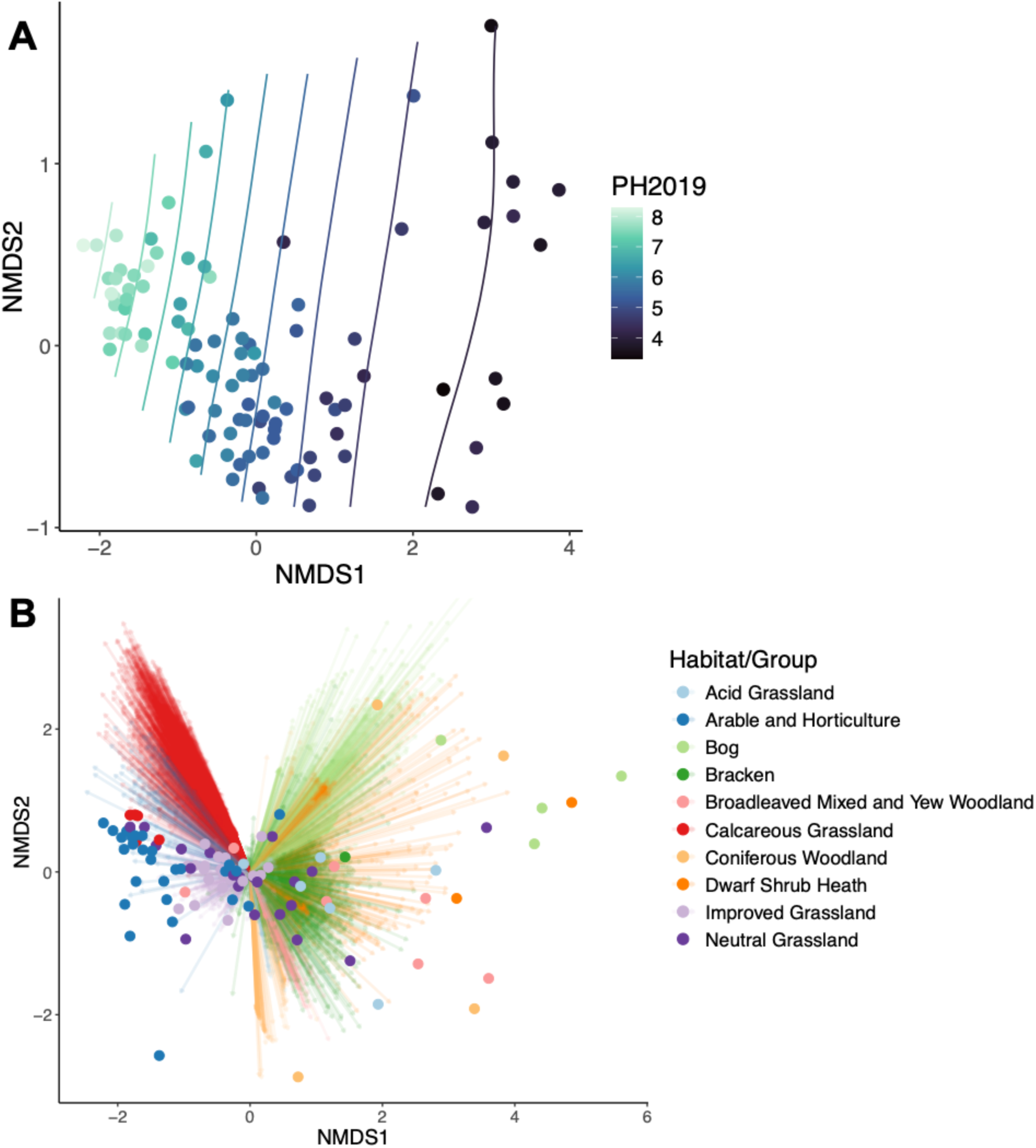
Ordinations of viral community composition. Variation in pH correlates with community dissimilarities (Bray-Curtis) (A). Lines denote the modelled pH surface of the ordination. Broad habitat class also shapes viral community composition (B). Points represent viral communities; arrows represent the species loadings of significantly associated vOTUs.

**Table 1.**
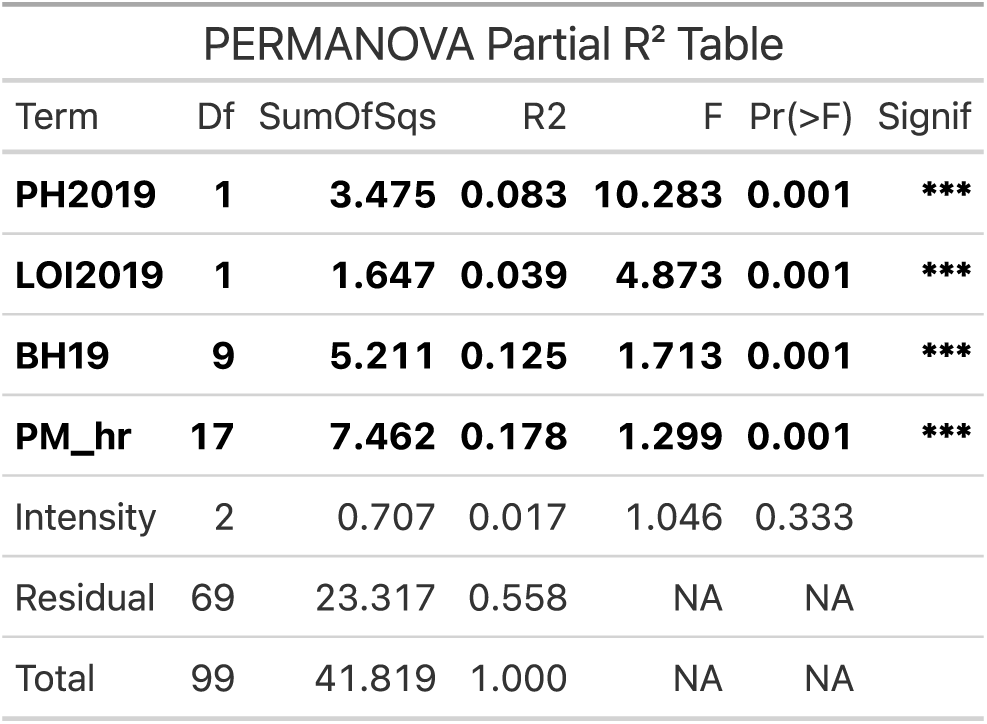
Drivers of viral community composition. PERMANOVA analysis of viral species community composition (Bray-Curtis distances) against environmental metadata with 999 permutations.

| PERMANOVA Partial R<sup>2</sup> Table |  |  |  |  |  |  |
| --- | --- | --- | --- | --- | --- | --- |
| Term | Df | SumOfSqs | R2 | F | Pr(>F) | Signif |
| **PH2019** | **1** | **3.475** | **0.083** | **10.283** | **0.001** | **\*\*\*** |
| **LOI2019** | **1** | **1.647** | **0.039** | **4.873** | **0.001** | **\*\*\*** |
| **BH19** | **9** | **5.211** | **0.125** | **1.713** | **0.001** | **\*\*\*** |
| **PM_hr** | **17** | **7.462** | **0.178** | **1.299** | **0.001** | **\*\*\*** |
| Intensity | 2 | 0.707 | 0.017 | 1.046 | 0.333 |  |
| Residual | 69 | 23.317 | 0.558 | NA | NA |  |
| Total | 99 | 41.819 | 1.000 | NA | NA |  |

To identify which viral sequences were driving these trends, and to assess the feasibility of using viral sequences as biomarkers, we ran an indicator species analysis. This analysis identifies species endemic to an environment that have both high specificity and fidelity. We typically found many sequences that were significantly associated with each of the broad habitat classes, with over 1000 viral sequences associated solely with calcareous grasslands and over 500 with bog environments (Fig 2B). By contrast, arable and horticultural soils and acid grasslands had fewer than 100. Neutral grasslands were unique in that they returned zero statistically significant markers; however, this category also has few diagnostic plant indicator species (Jackson, 2000), and may therefore aggregate multiple distinct environments into a catch-all term.

From these analyses, it was unclear whether broad habitat classes were meaningful for determining viral community composition, when they are partially confounded with pH, which is arguably more relevant to the local environment experienced by a virus and its host. We therefore focussed on improved and neutral grasslands, where the pH range of improved grasslands in nested within that of neutral grasslands. For this subset of samples, we still found significant effects of both pH (F_1,40_ = 3.5, p = 0.002) and habitat class (F_1,40_ = 1.96, p = 0.002). In this instance, pH explains more of the variation in viral community composition than habitat class, with a partial R^2^ of 8% vs. 4%. Taken together, our results suggest that pH is the major driver of viral ecology, but other environmental variables that are captured by the habitat categorisation also play a role in shaping patterns of viral biodiversity.

While environmental conditions will likely shape the viral community to some extent, a major determinant of viral communities must be the availability of hosts. We therefore predicted a strong correlation between viral community composition and bacterial community composition. To assess the extent that these populations are correlated, we used a 16S rRNA barcoding dataset collected from the same soil samples. Viral community dissimilarity (Bray-Curtis) correlated strongly with 16S bacterial community dissimilarity across all taxonomic ranks (Spearman Mantel tests, p < 0.0001). This correlation was strongest between viral and bacterial community at ASV resolution, (M = 0.901; Fig. 3). While correlation coefficients attenuated slightly at broader taxonomic ranks, strong associations were found at Genus (M = 0.695), Family (M = 0.792), Order (M = 0.845), Class (M = 0.765), and Phylum levels (M = 0.683). These correlations are consistent with previous reports of similar strength correlations using similar methods (Wicaksono et al. 2023).

**Fig. 3.**
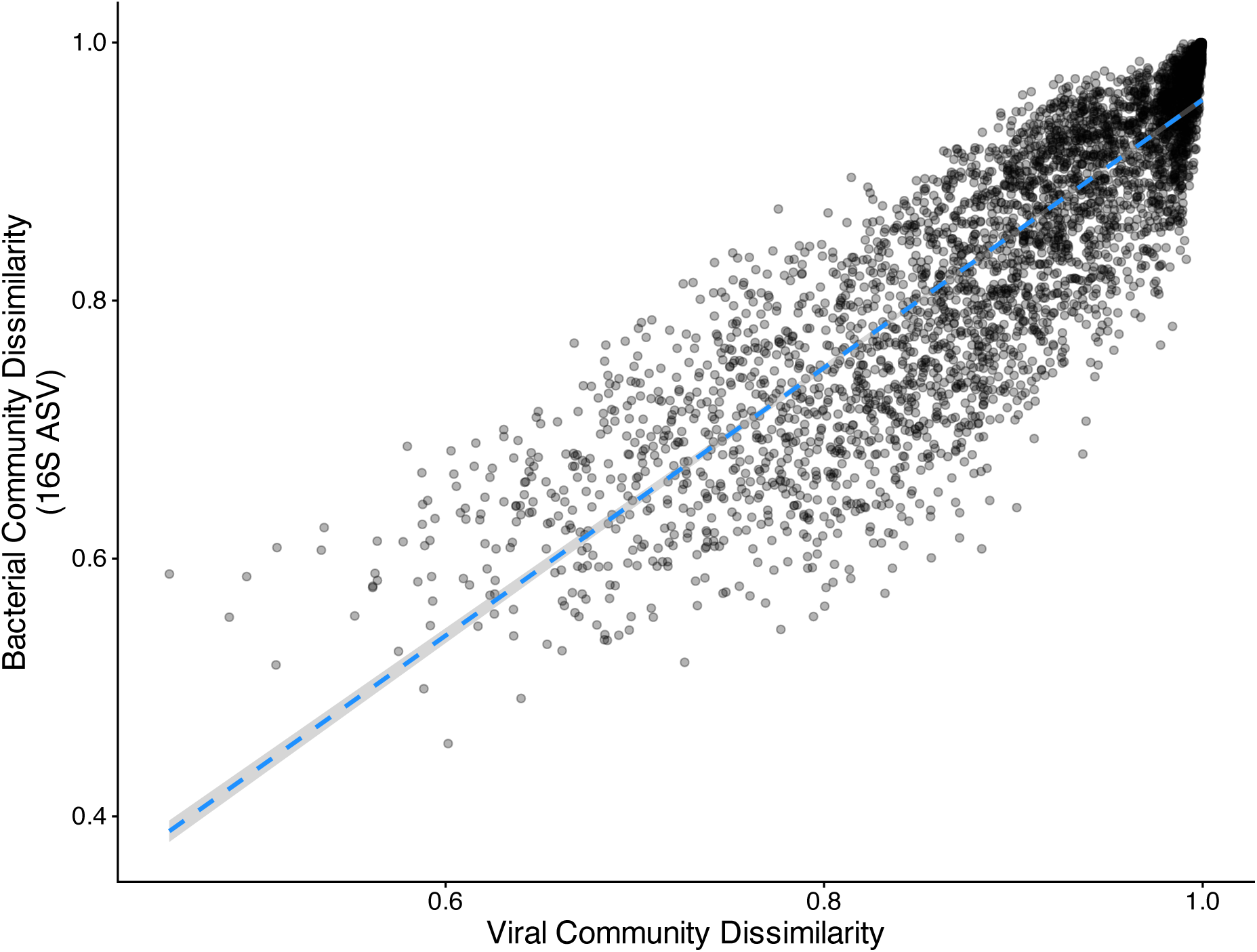
Viral community dissimilarity correlates with bacterial community dissimilarity. Points represent the pairwise dissimilarities between samples. Viral sequences are derived from metagenomic data while bacterial sequences are from an independent 16s rRNA barcoding dataset. Dashed line represents a linear model fit with shaded areas showing 95% confidence intervals. Mantel correlation = 0.901.

In sum, the strong environmental signal and indicator analysis demonstrated that many vOTUs must be common among samples of the same habitat. We investigated this further by asking how many sites each viral sequence was typically found in. Most viral contigs showed high spatial turnover, occupying a median of just 6 sites (IQR = 3 - 13), although there were exceptions, with one contig identified in 100% of samples, likely representing an integrated prophage in a widespread bacterial species (Fig. S5). When we restricted the analysis to genomes marked as medium, high quality or complete, we found slightly higher endemism, occupying a median of 4 sites (IQR = 2 - 7), with maximum occupancy from one genome that was identified in 65% of samples.

We then aimed to predict the putative hosts of the vOTUs, however the majority were too short to represent intact phage genomes. After quality assessment with CheckV we saw a sharp drop off in contig lengths between medium and low-quality genomes. While medium quality genomes were typically shorter than high-quality ones, they still averaged ∼39kb: within the bounds of a complete phage genome. Conversely low-quality genomes averaged 2.3kb and therefore represent fragments of viral genomes. We therefore extracted complete genomes along with high and medium quality phage genomes (n = 263 total) from the database (n = 58,545) and ran those through the host prediction tool IPhoP using the default database. Host prediction analysis returned few matches, with only 28 vOTUs (9 of which were identified as prophages) assigned to a host. The phages with the highest prevalence across sites (10 – 64%) were all predicted to infect members of the Pseudomonata, including Xanthobacteria, Acetobacteria, Bradyrhizobium and Beijerinckiaceae and these more prevalent vOTUs showed pH optima around pH 5.5 (Fig.4).

**Fig. 4.**
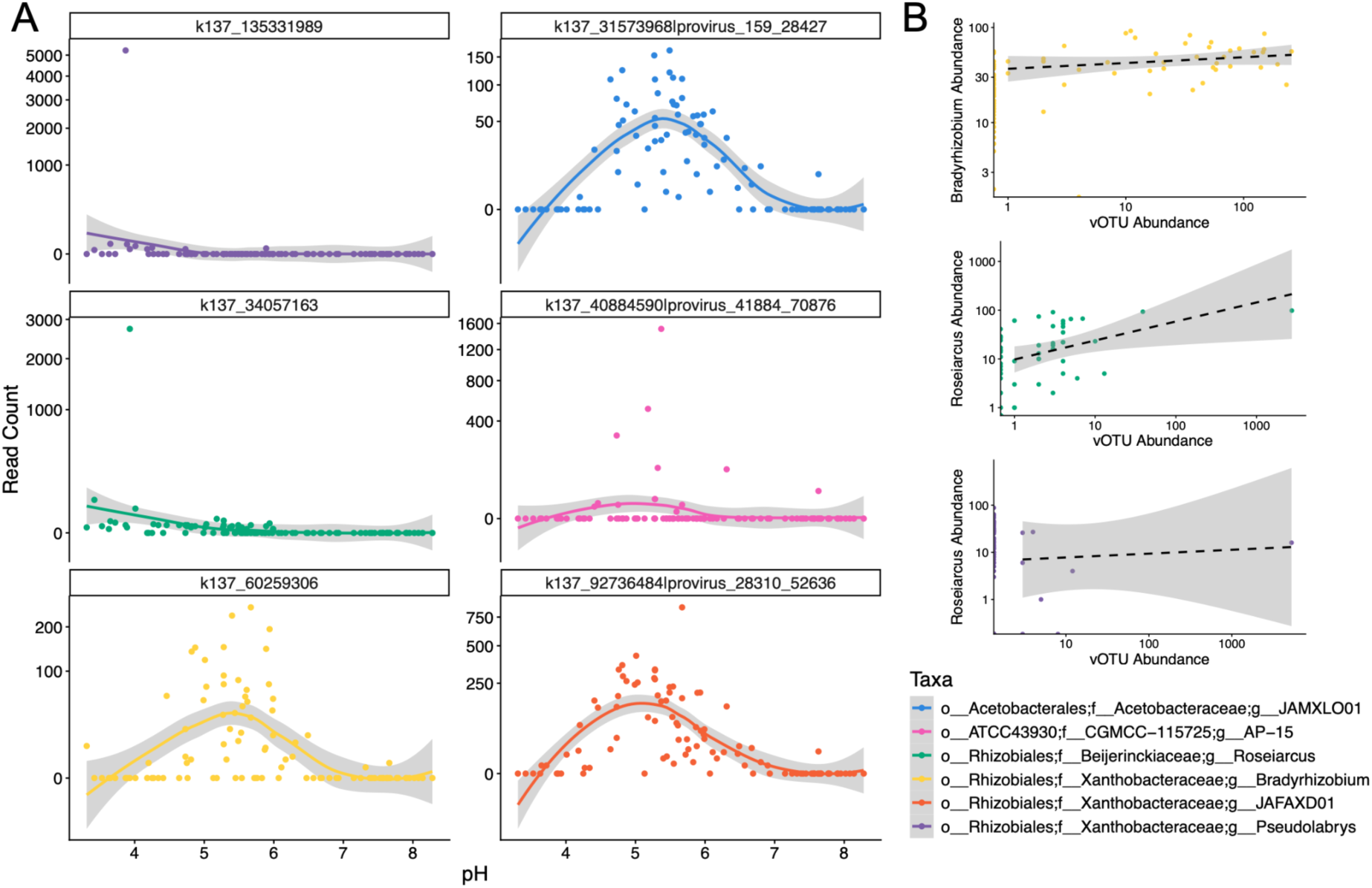
Common viral sequences are associated with weakly acidic soils. **(A)** Facets show the pH distributions of the most common 6 viral sequences. Solid line represents a generalized additive model fit with shaded areas showing 95% confidence intervals. (B) Correlations between a subset of vOTUs and their predicted hosts (Genus level).

### Virally encoded functional traits

We next explored the functional traits of these viral communities. Lysogeny is a major life-history trait of temperate phages, which we aimed to quantify across the survey. For this analysis, we linked integrase annotations, which we use as a proxy for a temperate lifestyle, to the abundance of viral contigs. While the presence of an integrase suggests potential for a temperate lifestyle it’s absence does not preclude it. Moreover, this analysis works at the contig level rather than a complete genome, which is required to predict a phage’s lysogenic potential. We found no significant effect of broad habitat class on the abundance of viral contigs containing KOfam-annotated integrase genes across the full dataset (F_9_,_90_ = 1.71, p = 0.09). This was also the case when we restricted the analysis to medium, high-quality, and complete genome subsets (F_9,90_ = 1.21, p = 0.30). Similarly, no habitat effect was observed when incorporating integrase annotations from GeNomad (F_9,90_ = 0.93, p = 0.50). Taken together, we conclude that habitat class does not determine frequencies of lysogeny, however we caution that integrase presence or absence is an imperfect proxy and further study with long-read datasets will be required to fully assess lysogeny rates.

We next assessed which, if any, virally-encoded functions were enriched in each habitat class. We performed differential abundance analysis for each environment against the average of the data (i.e., a one-vs-all analysis). No obvious trends emerged, but the full list of significantly enriched or depleted functions are available in the SI (Table S2). From this analysis, integrases were significantly underrepresented in a variety of habitats, spanning low pH environments like bog and bracken to the high pH calcareous grassland. Therefore, there is no obvious link between sample types and viral lifestyles in these data.

We also searched for AMR genes in the viral dataset, however we found no instances of these genes being carried on our viral contigs. We did however identify a number of beta-lactamase inhibitors, consistent with the idea that phages may represent a source of novel antimicrobial agents (Schroven et al. 2021).

#### Anti-phage defence systems

If viral communities are distinct among soil environments, a reasonable hypothesis is that they will select for different forms of bacterial immunity. These defence systems, such as CRISPR and RM systems, protect against invading phage and plasmid sequences via myriad modes of action. We searched for these defence systems in the full metagenomic assembly and assessed differences in abundance among environments. We found that the composition of defence systems significantly varied by habitat, LOI and pH (Table 2). The NMDS plots showed no obvious groupings of samples, yet the result was significant, suggesting some habitats select for specific defences, or for specific taxa with unique defence profiles. We therefore applied an indicator analysis, which assesses associations with environments based on specificity and fidelity (i.e., how exclusive a defence system is to a specific habitat, and how many of those sites within the habitat it is found). Overall, the majority of significant systems exhibited high fidelity but low sensitivity across samples, meaning they were generally widespread, a distinct subset demonstrated both high fidelity (1) and high sensitivity (>0.8). These were cas_type_V in bog environments (p < 0.05), cas_type_I-B1 in bracken (p < 0.05) and PDC-M71 in calcareous grasslands (p < 0.05). These results suggest that these defence systems may confer a selective advantage to bacterial hosts in these environments, or that the taxa that usually carry these defence systems are well-adapted to these niches.

**Table 2.** Environmental predictors of anti-phage defence composition.

| Term | Df | Sum Of Squares | Partial R2 | F | Pr(>F) |
| --- | --- | --- | --- | --- | --- |
| **Broad Habitat Class** | 9 | 0.389024977910522 | 0.13 | 3.04008038505746 | 0.001 |
| **pH** | 1 | 0.143214035882407 | 0.05 | 10.0724628356909 | 0.001 |
| **Loss on ignition** | 1 | 0.113325340790867 | 0.03 | 7.97034506027935 | 0.002 |
| **Residual** | 88 | 1.25121684370925 | 0.43 | NA | NA |
| **Total** | 99 | 2.84963398860569 | 1 | NA | NA |

## Discussion

Recent studies have developed global atlases of soil virus diversity (Graham et al, 2024, Ma et al. 2024, Goemann et al. 2026), but an understanding of landscape-level drivers of viral ecology is lacking for UK soils. We aimed to assess viral ecology by leveraging a metagenomic dataset collected as part of an ongoing nationwide monitoring campaign that used standardised collection methodologies. At the global level, traditional community ecology statistics are reported be unworkable, due to the extreme diversity present in the virosphere (Graham et al., 2024). However, in our dataset ordination analyses were tractable, indicating many shared sequences among samples of the same habitat types. Applying these ecological analyses, we found that multiple environmental factors were significant predictors of viral diversity (alpha and beta).

pH is a major driver of soil viral alpha and beta diversity in our dataset, somewhat consistent with previous experimental manipulations of a pH 4.5 – 7.5 gradient (Lee et al. 2022), although we saw a reduction in alpha diversity at pH 6.95 and above (arable and horticulture soils). Soil pH also shaped viral community composition, consistent with previous reports (Lee et al. 2022; Liao et al. 2022) which is intuitive, given the strong effects pH has on structural bacterial populations in soil both in UK soils (Griffiths et al. 2011) and globally (Bahram et al. 2018). Following this, our viral community composition was correlated with bacterial species composition. However, this could be due to a couple of reasons. Firstly, specific viruses track certain host taxa. Secondly, the same ecological pressures also select for specific phage types. These are challenging to disentangle and are not mutually exclusive. Evidence for the environment selecting for specific phage lifestyles comes from induction experiments, where lysogeny was more frequent during colder winter months (Jacoby et al. 2024). Understanding the contribution of environmental conditions that directly select for phage traits will be valuable future research.

Parent material, carbon content and broad habitat class were all also significant predictors of viral diversity. However, these results are correlational and many of these factors covary strongly. Therefore, future experimental manipulations of pH and carbon addition will be required to tease apart direct effects from covariance, ideally using soils collected from discrete broad habitat classes.

While soil virome experiments have demonstrated more extreme spatial turnover of viral sequences relative to bacterial hosts, we were unable to assess this due to the undisclosed nature of the sampling locations (Santos-Medellin et al. 2022). Our analysis of the paired samples, which compared proximally located high and low land use intensity sites within 1km^2^, found these pairs were as different from one another than to any other habitat. Therefore, factors relating to habitat and edaphic properties such as pH, are likely more strongly determining factors in community composition than geographic distance. We suggest that future efforts in UK soils assess distance-decay relationships of viral communities, to fully understand the dispersal limitations, if any, of viruses.

A limitation of this study and the approach used is that we only have information about the ‘bulk’ fraction of the metagenome, rather than the viral size fraction used for metaviromics. This likely skews the recovered sequences towards integrated prophages, and limits viral genome assembly due to dilution of sequences within a complex metagenome. Metaviromics, where the viral size fraction is sequenced, more accurately captures lytic, or recently lysed, virions (Santos-Medellin et al., 2021). Further, these data do not capture RNA or ssDNA viruses, but these also constitute the soil virome (Hillary et al. 2022). In the future, we suggest a collection of long-read bulk metagenomes paired with virome samples to generate a more precise picture of the soil virosphere. Emerging technologies, such as proximity-ligation sequencing and single-cell bacterial sequencing will allow more accurate host predictions, furthering our interpretation of infection patterns (Wu et al., 2023). Nonetheless, all samples here were collected using identical procedures, allowing fair comparisons to be made among samples, even if they represent a sub-section of the full virosphere.

The functional impacts of viruses in soils can range from beneficial to parasitic, depending on factors including cell-lysis rates, encoding of auxiliary metabolic functions and providing protection against competing viruses (Jansson and Wu, 2023). However, rates of lysogeny may vary between soils and there is likely be strong selection against prolonged extracellular exposure in harsh environments, as suggested by challenges with virome recovery from sulfuric soils (Bi et al. 2024). The roles of viruses in mediating soil carbon have been extensively reviewed, but broadly mediate carbon flux through cell lysis, and contribute to the soil carbon pool through viral biomass directly (Zimmerman et al. 2024). Exploring variation in these roles is therefore valuable for interpreting viral effects in UK soils. We used the abundance of a key marker of a temperate lifestyle, integrases, to assess the prevalence of this lifestyle. We found no significant effect of habitat class, but reiterate that these data are fragmented and largely composed of short sequences. Therefore, lifestyle prediction tools, such as BACPHLIP (Hockenberry and Wilke, 2021), will not accurately perform on these data. Future long-read datasets will allow more accurate assessment of lifestyles, as has been recently demonstrated in the human gut virome (Wirbel et al. 2026). Other functional trait analysis was largely uninformative, although some traits were significantly enriched in different habitats (SI). While viruses can encode auxiliary metabolic functions to confer host fitness benefits, recent debate around their identification suggests identified genes should be flanked by viral marker genes (Martin et al. 2025). The fragmented nature of our dataset means caution is required when interpreting AMGs, however we did identify, as an example, a beta-lactamase inhibitor encoded by a phage (Fig. 5). This demonstrates the utility of metagenomic analysis for identifying novel genes with biotechnology potential. We suggest that integrating long-read technology and viromics will yield valuable novel genetic substrates that could be exploited for various uses (Kosmopoulos and Anantharaman, 2025). Anti-phage defence systems, such as CRISPR, are also valuable for biotechnology and our analysis found some of these systems are enriched in specific environments. For example, Cas type V systems were enriched in bog samples. Future work might use similar approaches to identify environmental hotspots in which to search for novel effectors.

**Fig. 5.**
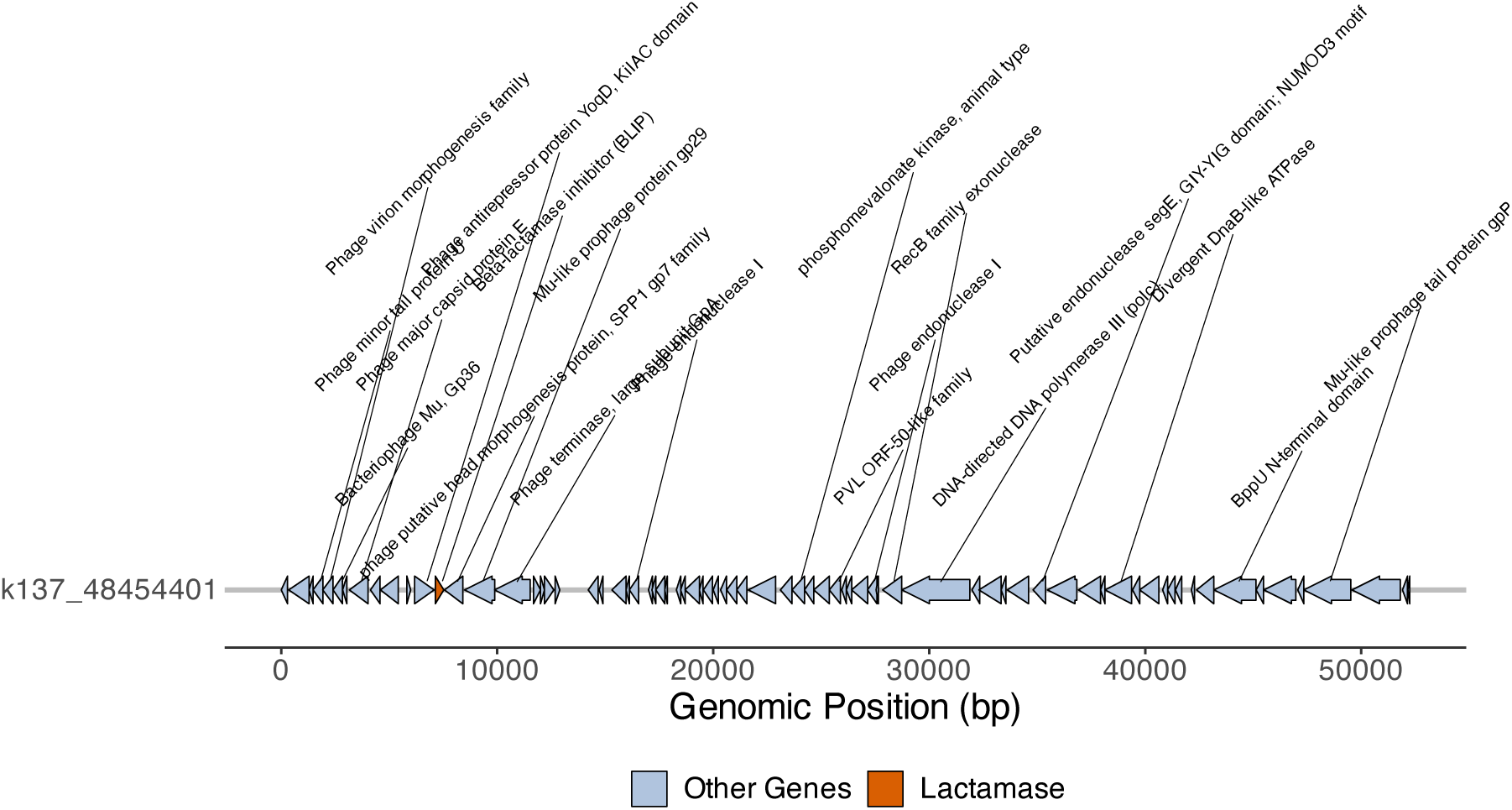
Soil phages as sources of novel antimicrobial candidates. Schematic of a complete phage genome carrying a Beta-lactamase inhibitor (orange), flanked by viral marker genes. Arrows represent predicted coding sequences with associated annotations from GeNomad.

While our results found many viral indicators species that are linked to the broad habitat classes, the majority of these vOTUs were rare. This is evident in the differences between species richness and Shannon index value, which suggests differences in species evenness. A long-tail of rare vOTUs in the rank-abundance curve shows many rare taxa. Our indicator analysis will discard any vOTU not deterministic of a land-use category, despite potential similarity in the immediate local soil environment. More generally, the functional importance of these rare vOTUs is unknown. The following question remains: do they represent viruses of rare, but active, microbes or are they akin to a viral seed bank, awaiting the blooming of a suitable host. Experiments from Mediterranean grassland soils suggests the latter, with massive viral biomass increases following water addition and subsequent host activity preceding viral lysis (Nicolas et al. 2023).

This characterisation of UK soil viral diversity gives insight into the understudied soil virosphere. Emerging molecular and bioinformatic tools will further our understanding of viral community ecology in terrestrial environments, and experimental manipulations will determine the functional role of viruses in soils. Incorporating these measurements into long-term monitoring programmes, such as the Countryside Survey, will allow us to assess viral dynamics alongside broader trends in soil health.

## Acknowledgements

This work was supported by a BBSRC fellowship (BB/X009793/1 to SM) and Natural Environment Research Council award number NE/R016429/1 as part of the UK-SCaPE programme.

## CRediT author statement

SM: Conceptualization, Formal analysis, Investigation, Writing – Original Draft, Visualization, Funding acquisition. BJ: Formal Analysis, Writing – Review & Editing. AS: Formal analysis. TG: Investigation. SBB: Investigation, Writing – Review & Editing. RG: Investigation, Writing – Original Draft.

## Notes

### Competing Interest Statement

The authors have declared no competing interest.

