## Supplementary Information for "Environmental drivers of soil virus diversity across UK habitats"

### Supplementary Info

Table S1. Statistical support for Fig.S1. Post-hoc testing of read depth vs. broad habitat class.

| contrast | estimate | SE | df | t.ratio | p.value |
|---|---|---|---|---|---|
| Acid Grassland - Arable and Horticulture | -1993192 | 4090000 | 90 | -0.487 | 1.0000 |
| Acid Grassland - Bog | -550406 | 5870000 | 90 | -0.094 | 1.0000 |
| Acid Grassland - Bracken | 5406700 | 9820000 | 90 | 0.551 | 0.9999 |
| Acid Grassland - Broadleaved Mixed and Yew Woodland | 3703208 | 5060000 | 90 | 0.732 | 0.9992 |
| Acid Grassland - Calcareous Grassland | 2607759 | 5510000 | 90 | 0.474 | 1.0000 |
| Acid Grassland - Coniferous Woodland | 970725 | 5870000 | 90 | 0.165 | 1.0000 |
| Acid Grassland - Dwarf Shrub Heath | 16085960 | 7420000 | 90 | 2.167 | 0.4867 |
| Acid Grassland - Improved Grassland | -2509526 | 4190000 | 90 | -0.599 | 0.9998 |
| Acid Grassland - Neutral Grassland | 494722 | 4210000 | 90 | 0.118 | 1.0000 |
| Arable and Horticulture - Bog | 1442786 | 4860000 | 90 | 0.297 | 1.0000 |
| Arable and Horticulture - Bracken | 7399892 | 9250000 | 90 | 0.800 | 0.9984 |
| Arable and Horticulture - Broadleaved Mixed and Yew Woodland | 5696400 | 3840000 | 90 | 1.483 | 0.8955 |
| Arable and Horticulture - Calcareous Grassland | 4600951 | 4410000 | 90 | 1.042 | 0.9887 |
| Arable and Horticulture - Coniferous Woodland | 2963917 | 4860000 | 90 | 0.610 | 0.9998 |
| Arable and Horticulture - Dwarf Shrub Heath | 18079152 | 6650000 | 90 | 2.717 | 0.1829 |
| Arable and Horticulture - Improved Grassland | -516334 | 2590000 | 90 | -0.199 | 1.0000 |
| Arable and Horticulture - Neutral Grassland | 2487914 | 2620000 | 90 | 0.948 | 0.9943 |
| Bog - Bracken | 5957106 | 10200000 | 90 | 0.586 | 0.9999 |
| Bog - Broadleaved Mixed and Yew Woodland | 4253614 | 5700000 | 90 | 0.746 | 0.9991 |
| Bog - Calcareous Grassland | 3158166 | 6100000 | 90 | 0.518 | 1.0000 |
| Bog - Coniferous Woodland | 1521131 | 6430000 | 90 | 0.237 | 1.0000 |
| Bog - Dwarf Shrub Heath | 16636367 | 7870000 | 90 | 2.113 | 0.5236 |
| Bog - Improved Grassland | -1959120 | 4940000 | 90 | -0.396 | 1.0000 |
| Bog - Neutral Grassland | 1045129 | 4960000 | 90 | 0.211 | 1.0000 |
| Bracken - Broadleaved Mixed and Yew Woodland | -1703492 | 9720000 | 90 | -0.175 | 1.0000 |
| Bracken - Calcareous Grassland | -2798940 | 9960000 | 90 | -0.281 | 1.0000 |
| Bracken - Coniferous Woodland | -4435975 | 10200000 | 90 | -0.436 | 1.0000 |
| Bracken - Dwarf Shrub Heath | 10679261 | 11100000 | 90 | 0.959 | 0.9938 |
| Bracken - Improved Grassland | -7916226 | 9300000 | 90 | -0.852 | 0.9974 |
| Bracken - Neutral Grassland | -4911977 | 9310000 | 90 | -0.528 | 0.9999 |
| Broadleaved Mixed and Yew Woodland - Calcareous Grassland | -1095449 | 5320000 | 90 | -0.206 | 1.0000 |
| Broadleaved Mixed and Yew Woodland - Coniferous Woodland | -2732483 | 5700000 | 90 | -0.480 | 1.0000 |
| Broadleaved Mixed and Yew Woodland - Dwarf Shrub Heath | 12382752 | 7290000 | 90 | 1.699 | 0.7933 |
| Broadleaved Mixed and Yew Woodland - Improved Grassland | -6212734 | 3950000 | 90 | -1.575 | 0.8567 |
| Broadleaved Mixed and Yew Woodland - Neutral Grassland | -3208485 | 3970000 | 90 | -0.809 | 0.9983 |
| Calcareous Grassland - Coniferous Woodland | -1637034 | 6100000 | 90 | -0.268 | 1.0000 |
| Calcareous Grassland - Dwarf Shrub Heath | 13478201 | 7610000 | 90 | 1.772 | 0.7507 |
| Calcareous Grassland - Improved Grassland | -5117286 | 4500000 | 90 | -1.136 | 0.9795 |
| Calcareous Grassland - Neutral Grassland | -2113037 | 4520000 | 90 | -0.467 | 1.0000 |
| Coniferous Woodland - Dwarf Shrub Heath | 15115235 | 7870000 | 90 | 1.920 | 0.6559 |
| Coniferous Woodland - Improved Grassland | -3480251 | 4940000 | 90 | -0.704 | 0.9994 |
| Coniferous Woodland - Neutral Grassland | -476002 | 4960000 | 90 | -0.096 | 1.0000 |
| Dwarf Shrub Heath - Improved Grassland | -18595487 | 6710000 | 90 | -2.769 | 0.1631 |
| Dwarf Shrub Heath - Neutral Grassland | -15591238 | 6730000 | 90 | -2.317 | 0.3884 |
| Improved Grassland - Neutral Grassland | 3004249 | 2770000 | 90 | 1.083 | 0.9852 |

P value adjustment: Tukey method for comparing a family of 10 estimates

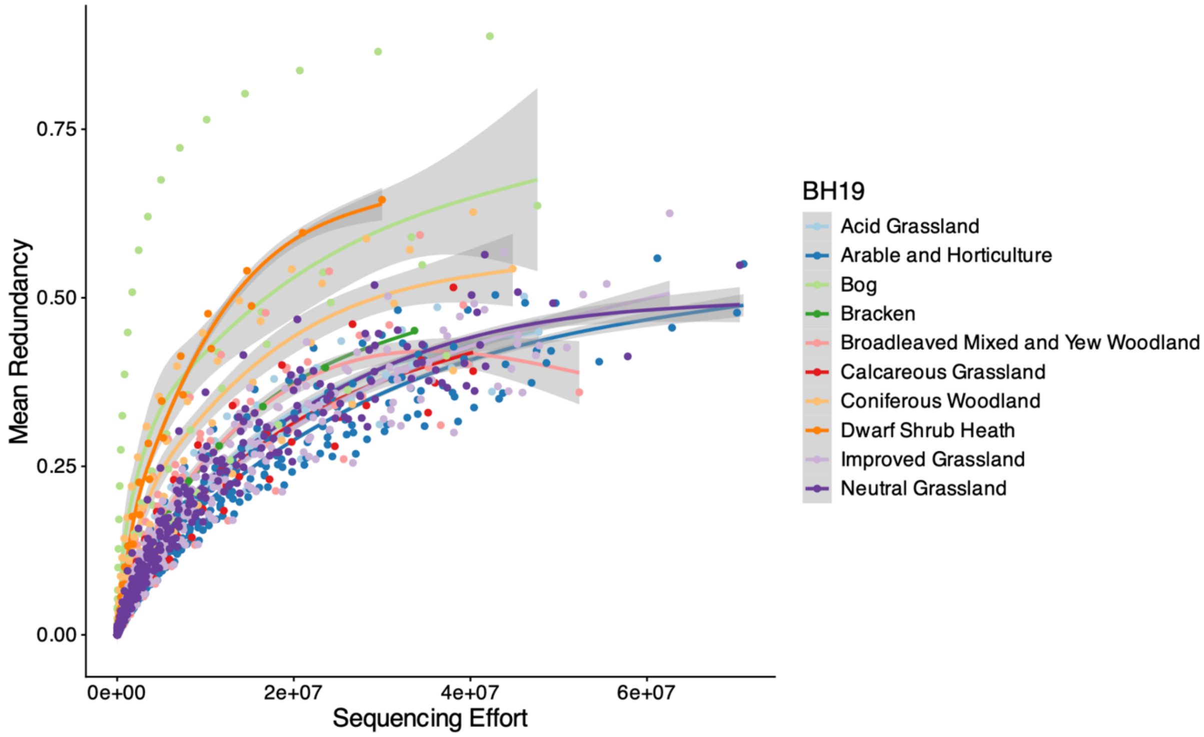


**Fig. S1. Coverage redundancy curves across sampling effort.** Lines show the mean redundancy of each group (n = 100 samples in total). Points represent individual samples, with 100 subsamples at each level of sequencing effort being used to estimate redundancy per biological sample. Data generated with Nonpareil3.

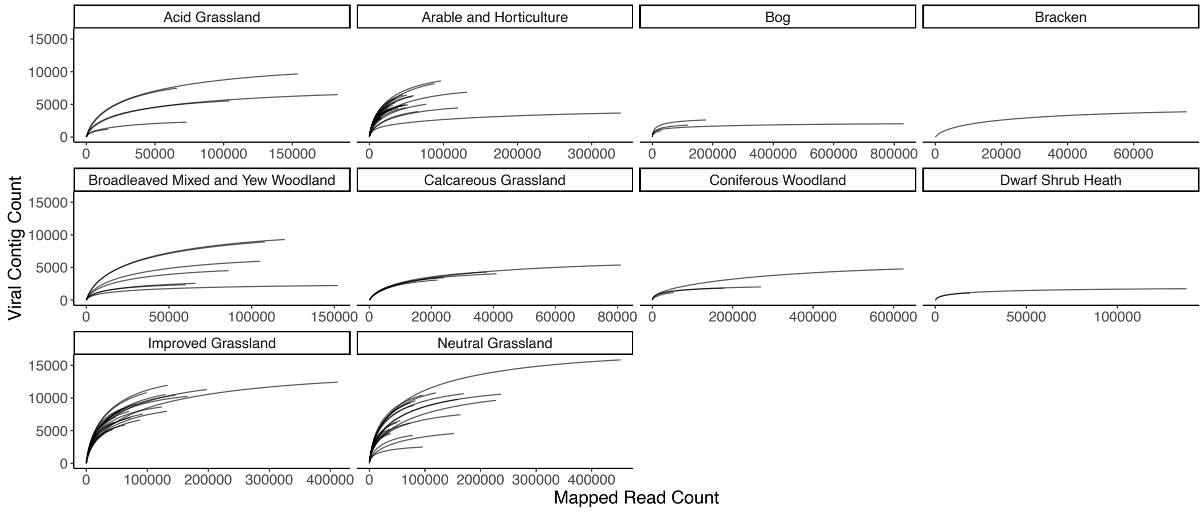


**Fig. S2. Rarefaction curves from the relaxed database split by sample.** Curves represent number of viral contigs that recruit reads at the various read mapping depths. Note that the asymptote represents saturation within our assembly, rather than the total environmental viral diversity.

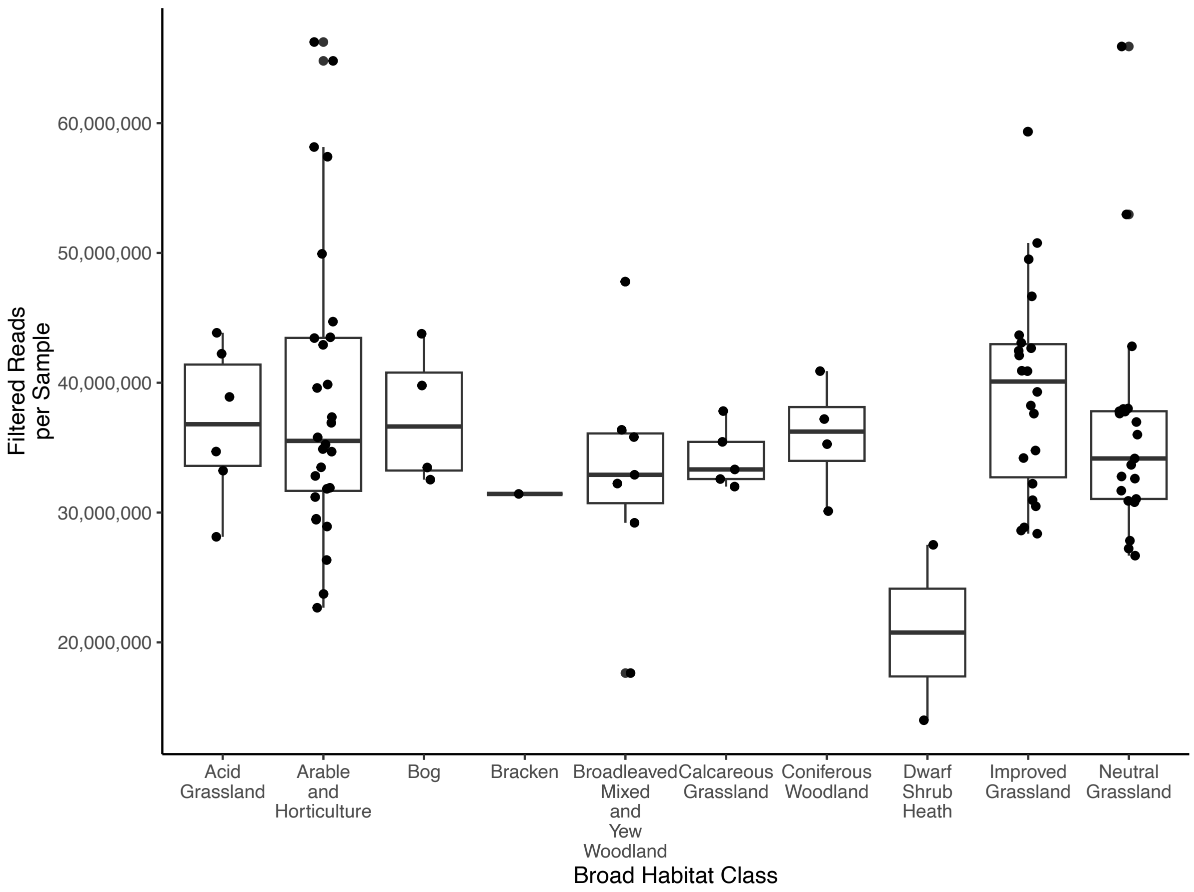


**Fig. S3. Read counts per sample grouped by Broad Habitat class.** Points represent the number of R1 reads per sample after quality filtering.

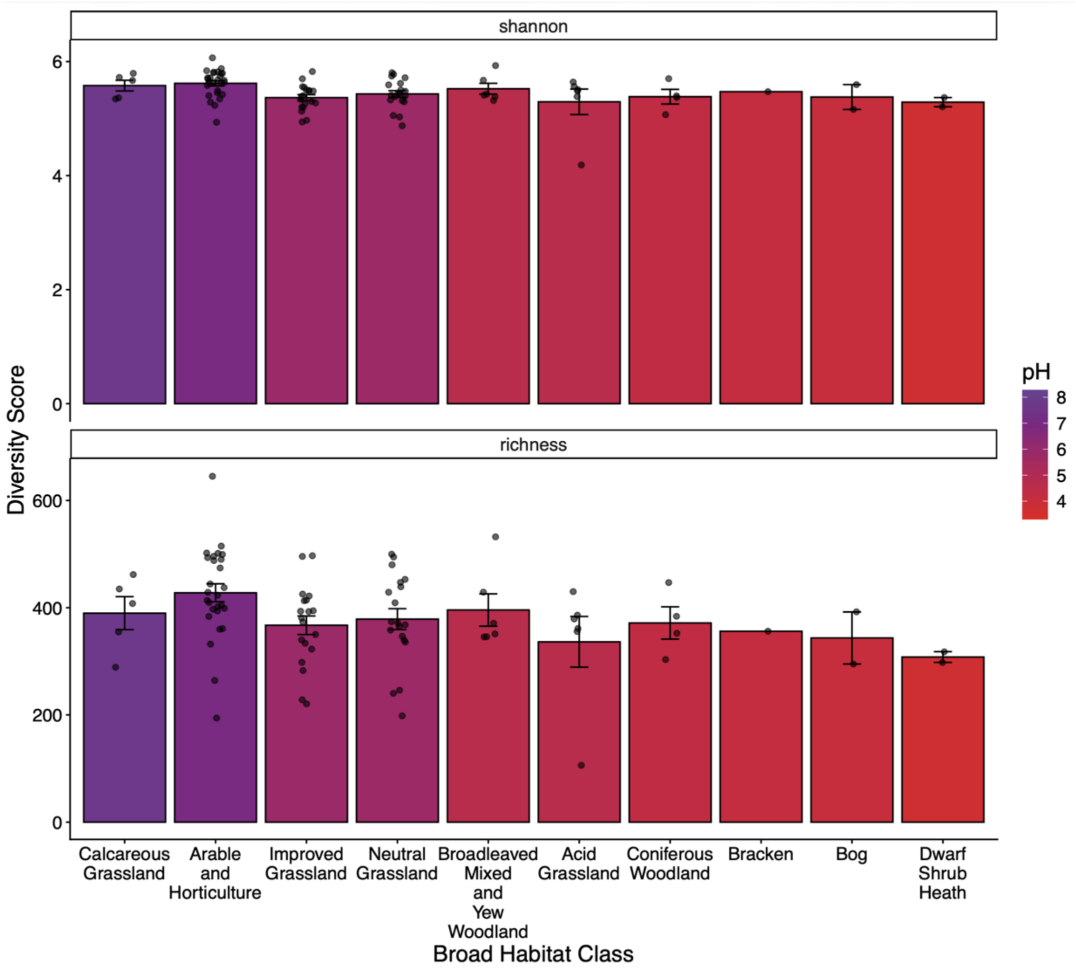


**Fig. S4. Bacterial ASV alpha diversity varies among habitat classes and with pH.** ASVs were generated from a parallel 16S rRNA experiment using the same soil cores. Data were processed independently as described in Goodall et al. 2026.

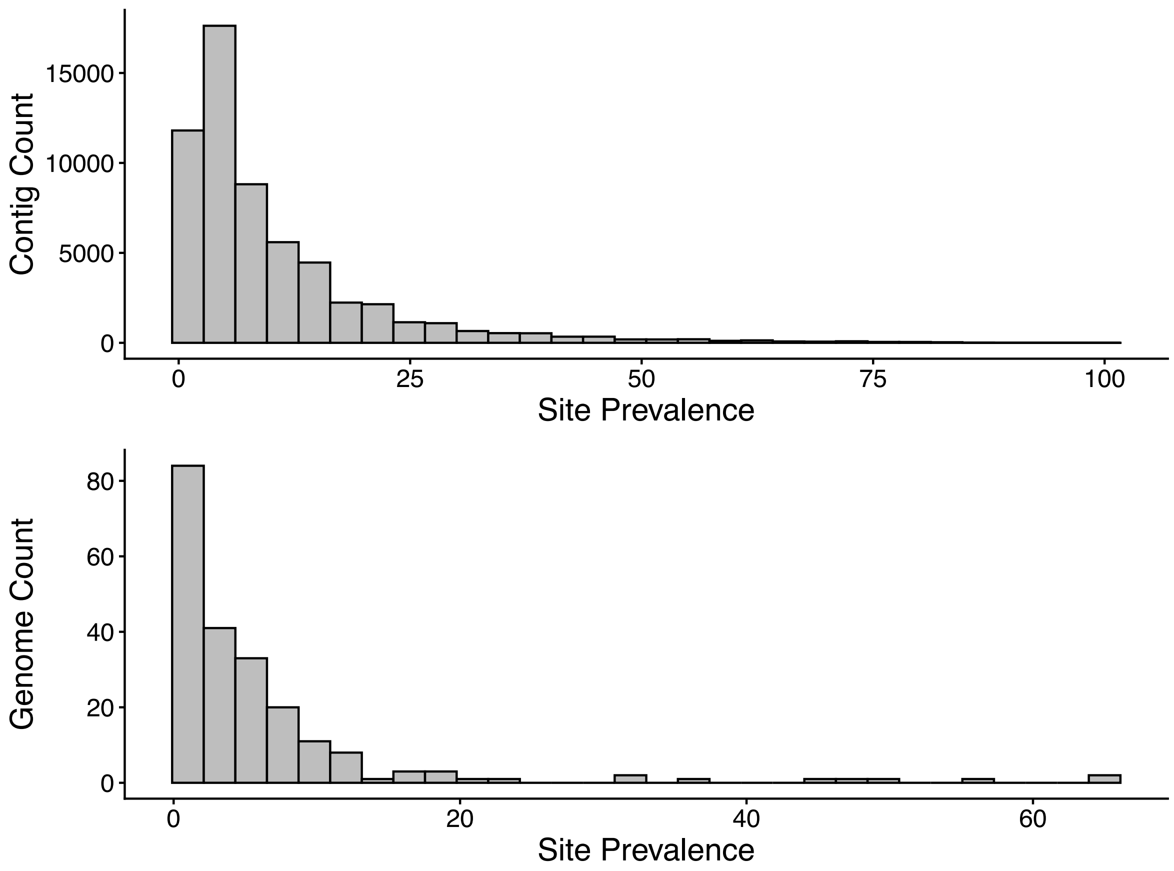


**Fig. S5. Site prevalence of identified viral sequences is low.** Most viral contigs (top) and genomes (bottom; based on estimated completeness as medium or higher) occur in few sites. Histograms derived from vOTU tables from the 'relaxed' database.
